# A harmonized ovarian cancer scRNA-seq atlas to dissect disease heterogeneity underlying metastatization and chemoresistance

**DOI:** 10.1101/2025.02.11.637633

**Authors:** Marta Rosa Sallese, Vittorio Aiello, Carlo Emanuele Villa, Pietro Lo Riso, Giuseppe Testa

**Affiliations:** Department of Oncology and Hemato-Oncology, University of Milan, Milan, Italy; Department of Experimental Oncology, European Institute of Oncology IRCCS, Milan, Italy; Human Technopole, Milan, Italy

## Abstract

The advent of single cell technology has enabled researchers with the ability to achieve unprecedented resolution in the characterization of biological systems. The consequent increasing availability of single cell dataset brought about the possibility to obtain comprehensive reference atlases, to shed light on the molecular and cellular foundations of healthy and diseased tissues. This process has highlighted the need to integrate datasets from different sources while being able to distinguish true biological signal from technical confounders. While this issue is widespread to most biological settings, it holds especially true for cancer samples, which are characterized by a diffused inter- and intra-patient phenotypic heterogeneity. To address this issue, here we developed a novel integration method tailored to highly heterogeneous single cell transcriptomic data and applied it to one of the quintessential heterogeneous cancer type, namely high-grade serous ovarian cancer, to generate the first reference atlas for this disease. By identifying patient-specific cell populations and deriving metacells, we were able to preserve inter-patient biological variability. Using a variational autoencoder, we integrated metacell data, revealing an evolving landscape of cell states along disease progression and treatment for each of the main cell types constituting the dataset. Also, we showed the potential of this resource by identifying diffused and tissue/treatment-specific cell-to-cell interactions. Finally, the generated integration model allowed to expand the atlas with additional data, granting iterative refinement over time of this disease reference. Our strategy now provides a valuable resource for the cancer research community, facilitating the investigation of tumor heterogeneity towards the development of novel therapeutic strategies.

## 1 Introduction

Cancer is a complex and dynamic disease, continuously shaped along tumor evolution, and thus characterized by a remarkable heterogeneity. Among the processes leading to such heterogeneity, the frequent genomic instability leads to the progressive accumulation of genetic and epigenetic aberrations and, thus, to the emergence of different subclonal populations carrying specific characteristics [1]. This autonomous effect is contrasted during the course of disease progression by both environmental pressure, due to the dynamic equilibrium established between the tumor and its surrounding microenvironment (TME), and by therapeutical challenge, leading to the emergence or selection of therapy-resistant clones [2]. Also, the capability to metastatize to distal locations directs tumor cells towards a further evolutionary challenge caused by the change in surrounding tissue and, consequently, in the tumor-TME interaction, leading to a disease that is very different from the primary one [3]. The advent of single cell technologies has overcome the limitations of the bulk profiling and are now allowing the investigation of such extraordinary heterogeneity and complexity at an unprecedented degree of resolution. In particular, the deluge of single cell transcriptomic data now allows the generation of atlases that serve as reference for the investigation of heterogenous and complex diseases [4] by providing a map of cell states linked to a variety of clinical and biological metadata and potentially hinting at the identification of clinically-relevant disease-associated phenotypes [5]. The intrinsic challenges in the generation of atlases are mainly related to single-cell technology susceptibility to technical batch effects which can arise when aggregating datasets generated in different laboratories, at different times and by different operators, and to the consequent need to correct data according to such batches while preserving biological information describing the data (e.g. the specific mutational burden characterizing each patient). Additionally, specifically for cancer and in contrast with the generation of healthy tissues atlases, the complexity of the disease and the absence of a priori knowledge of the existing cell subpopulations poses a significant challenge for the definition of a coherent and informative atlas. Indeed, in this context there is the need to disentangle the several layers of biological heterogeneity intrinsically bound to the disease from the technical data heterogeneity. Beside these, there is the problem related to the harmonization of such a huge amount of data, in terms of collecting the same categories of metadata and assigning them univocal descriptions. For this reason, we developed a framework called Single Cell transcriptomic Atlas Integration Pipeline (SCAIP) (Fig. 1). Here we aimed at building a comprehensive resource in the form of a single cell transcriptomics atlas and related data integration strategy for the investigation of disease heterogeneity. We used as a paradigmatic example of heterogeneous cancer disease the most lethal gynecological malignancy, High-Grade Serous Tubo-Ovarian Cancer (HGSTOC). The characterization of HGSTOC tumor biology at single cell resolution is currently hampered by the lack of studies with a sufficient number of tumor cells and patients, both due to the high prevalence of non-tumoral cell types and the financial burden of conducting such experiments at scale. Thus, our atlas tackles this need by aggregating together 9 different publicly available datasets and developing a related data integration strategy able to preserve biological heterogeneity (Fig. 1). To obtain a faithful representation of data heterogeneity and describe the source of this heterogeneity we performed clustering and metacells derivation separately for every sample to identify the distinct subpopulations characterizing each patient. In addition, to robustly represent the space describing the ecosystem of tumor and tumor-associated cell populations, i.e. cancer cells, and cancer-associated immune (CAI), fibroblast (CAF) and endothelial (CAE) cells, for each of the latter we selected the space identified by the union of highly variable genes, computed separately on every patient, to preserve the variability of the system. Then, we took advantage of a deep learning-based data integration tool, namely a variational autoencoder, and we fed it with metacells data to integrate and characterize the different subpopulations that may be similar across samples. After integration, we were able to identify new robust gene signatures that would have been masked by datasets heterogeneity. Moreover, we have been able to identify tissue type- and treatment-specific transcriptomic features and to retrospectively map them on the un-integrated datasets, also increasing the statistical power to enable the identification of disease-relevant phenotypes in newly generated experiments. This deep-learning integration strategy provides a robust framework to easily and iteratively expand the atlas with newly generated datasets, thus constituting also an invaluable resource for the scientific community tackling HGSTOC pathogenetic mechanisms. Moreover, while we used HGSTOC as a proof of concept, the underlying rationale of our feature selection approach represents a concept that could potentially be applied to other heterogeneous diseases.

**Fig.1:**
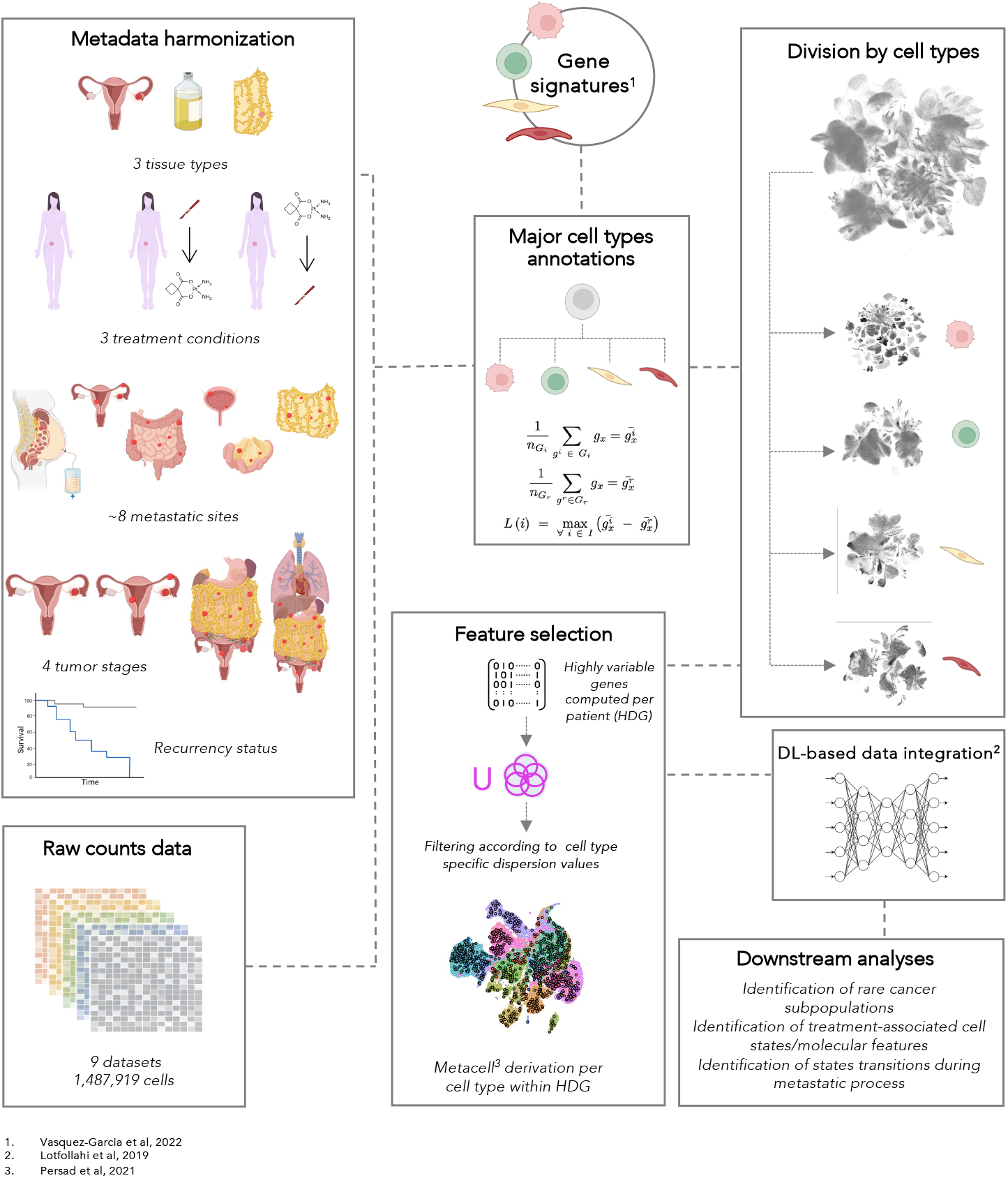
Overview of the HGSOC Single cell transcriptomic Cancer Atlas Integration Pipeline (SCAIP). Harmonized metadata and raw counts were labelled using predefined gene signatures to distinguish the four major cell types characterizing the tumors. This labels were used to divide the atlas into four cell subsets. For each of them we separately computed highly defining genes (HDG) and derived metacells within this feature space. Then each metacells subset was integrated using a deep learning (DL) based integration strategy. Subsequently these features were used to perform downstream analysis aimed at tumor characterization and investugation of its interaction with tumor microenvironment (TME).

## 2 Results

### 2.1 OvCA reveals different degrees of heterogeneity across major populations

To build the Ovarian Cancer Atlas (OvCA) we collected 9 scRNA-seq datasets [6]–[14] including associated clinical metadata for a total of 1.5M of annotated cells. To avoid variability related to different sequencing technologies, we only selected datasets with cells profiled by 10X Genomics technology. These datasets include data from 79 different patients, 3 different tumor tissue types (primary ovarian site surgical specimens, ascitic fluid and metastatic tissue specimens). Specifically for metastatic samples, we also aggregated information about the anatomical site of the metastasis and found the omentum and peritoneum the most common sites from which samples were retrieved, as expected [15] (Fig. 2A-B). The dataset entails also indication of the therapeutic status of patients, including naïve patients (i.e. pre-chemotherapy samples) and patients who received a treatment according to the most common therapeutic regimens, i.e. either following post-debulking adjuvant chemotherapy (CHT) or neoadjuvant chemotherapy (NACT) prior to de-bulking surgery. Where available, we also reported recurrency status and tumor stage (Fig. 2A, Supplementary Fig. S1A, B). The majority of samples were obtained from patients with metastatic disease (68 out of 79 samples), which is the most common stage of presentation for this cancer [16] given the high prevalence of late stage diagnosis high grade serous ovarian tumors [16]. Accordingly, we found the vast majority of samples being stage III or IV according to FIGO staging.

**Fig.2:**
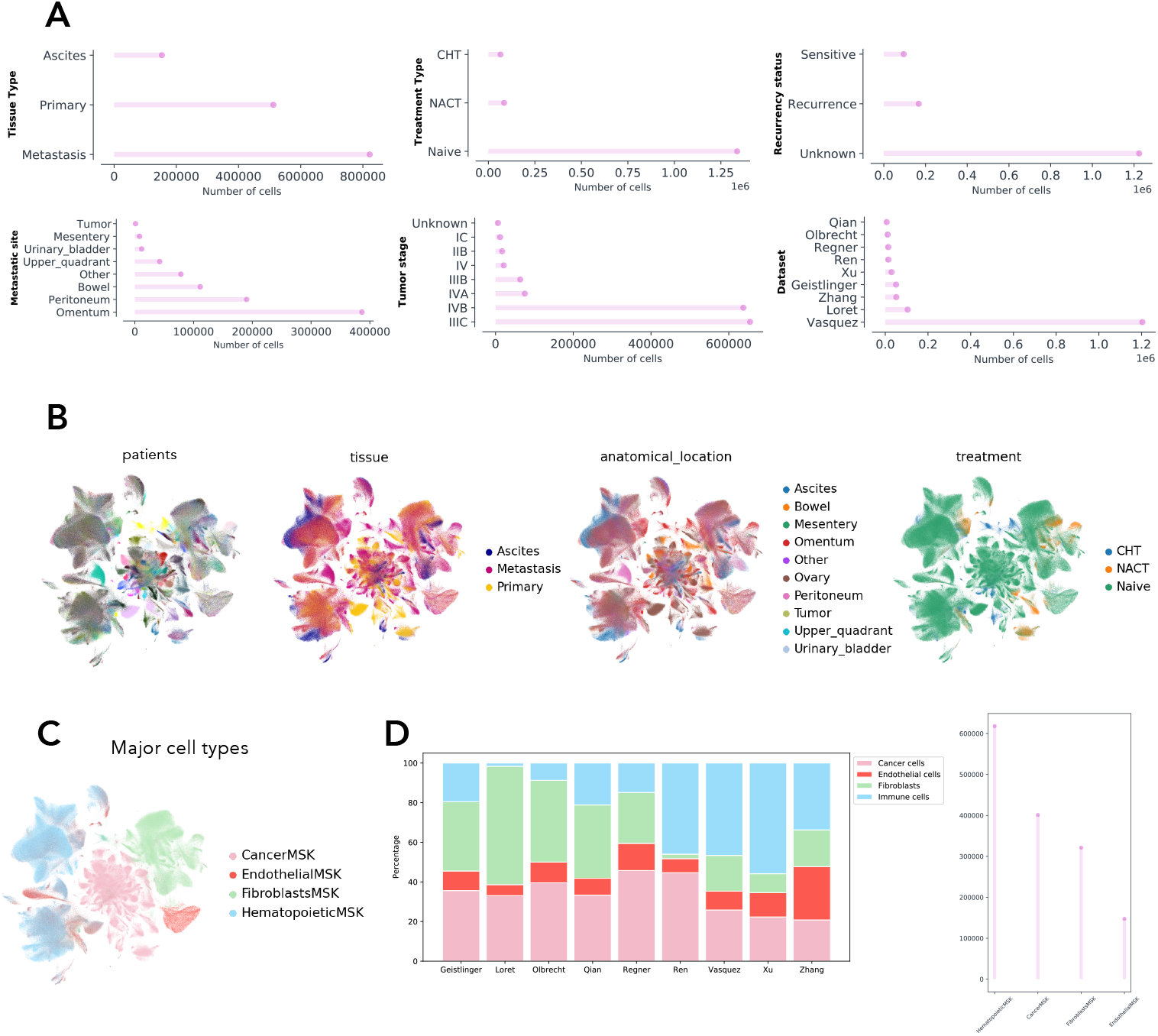
OvCA reveals different degrees of heterogeneity across major populations. A. Patients and samples composition of the Ovarian Cancer Atlas (OvCA) displayed by tissue types from which samples derived, treatment that patients’ undergone (chemotherapy named CHT and neoadjuvant chemotherapy named NACT), recurrency status, metastatic site (here OvCA was subsetted to only show metastatic samples), tumor stage (according to FIGO classification) and dataset from which they were provided. B. UMAPs displaying OvCA colored by (in order from left) patients (being 79 patients here we are not showing legend because of space constraints), tissue types, metastatic site (primary tumors (named here ‘Ovary’) and ascites are displayed too) and treatment. C. UMAP displaying the four major cell types populations defined by literature-based signatures. D. Percentages (on the left) and absolute numbers (on the right) of each dataset composition in the four main cell types we derived.

Given the well-known heterogeneity of HGSTOC tumors, we reasoned that a more meaningful and accurate representation of the data could be obtained by unpacking the complexity of our system into simpler and more coherent spaces in terms of expression variability. To this aim, we first proceeded with the identification of the major cell types characterizing the atlas that could then be used to define more homogenous spaces describing the tumor components. Using recently described marker gene signatures [11] we created an a priori cell type-labelling strategy to distinguish the major cell types of the dataset (Cancer cells, CAI, CAF and CAE cells). We derived four scores, one for each major cell type (see Methods), to identify cells pertaining to each subset. We scored 400,484 cancer cells, 640,262 CAI, 321,193 CAF and 125,980 CAE cells (Fig. 2C). Across datasets, the tumoral component of the samples was consistently lower than 40%. For the other components, fluctuations in the percentage of cells pertaining the specific cell type could be attributed to the processing procedure, for instance CD45+ cells depletion prior to sequencing such as in Loret et al., or to the representation of the tissue types in the dataset, such as in Ren et al. which entailed exclusively ascitic samples, with ensuing reduced representation of stromal cells. (Fig. 2D). Notably, despite observing a strong dataset imbalance in terms of the global number of cells derived by Vasquez-Garcia et al (Fig. 2A, Supplementary Fig. S1C), the overall heterogeneity was not affected and remained mainly due to the biological difference of the four major cell type populations. Then, we subset the atlas according to the four cell types and assessed for each of them the degree of inter-patient heterogeneity. We observed that cell types were different in terms of intra- and inter-patient heterogeneity. Cancer cells showed a considerably more scattered distribution of individual patients across the manifold compared to the other cell types (Fig. 3A) while immune cells were the most homogenous cell type (Fig. 3B). To quantify patient heterogeneity across the four major cell types we computed clustering quality metrics including Adjusted Rand Index (ARI) [17], Adjusted Mutual Information (AMI) [18], Fawlkes-Mallows Index (FMI) [19] (See Methods). Cancer cells scored considerably higher (Fig. 3C) than CAF, CAE and CAI cells (Fig. 3D demonstrating that inter-patient variability affects clustering mainly in cancer cells, while in the other cell types, clustering is affected by other sources of variability. These strong differences among cell types warrant it as more meaningful to treat them separately as coherent systems for the downstream procedures, consistent with them being distinct biological objects with different sets of features lying in different manifolds. In line with this reasoning, in the first analytical steps each cell type was subjected separately to the tasks described below.

**Fig.3:**
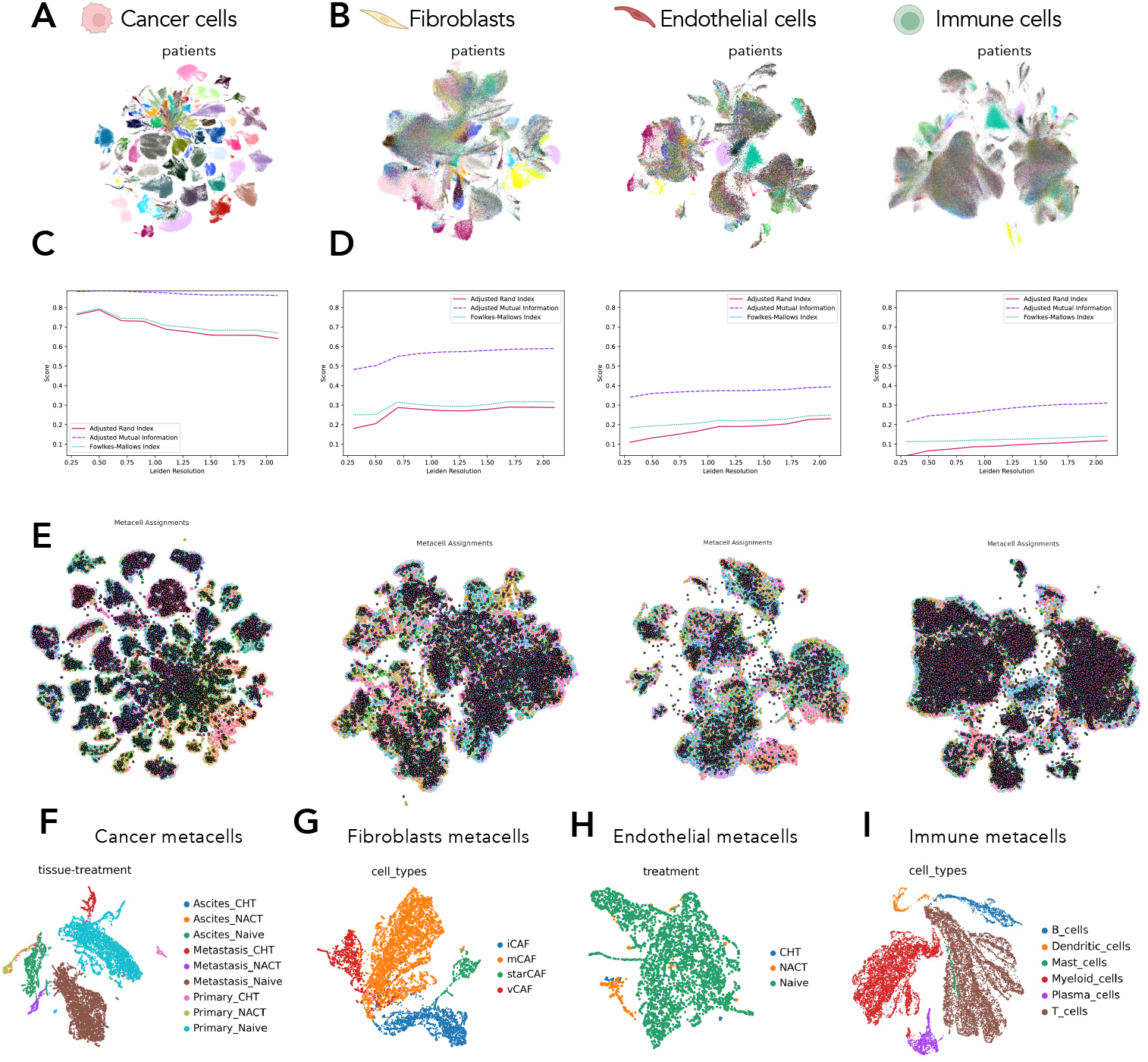
Data integration allowed to characterize cancer, endothelial and fibroblasts cell states across tissues. A. UMAP displaying cancer cells from OvCA colored by patients (being 79 patients here we are not showing legend because of space constraints). B. UMAPs displaying cancer-associated (from left) fibroblasts, endothelial and immune cells from OvCA colored by patients (being 79 patients here we are not showing legend because of space constraints). C-D. Overall scores for ARI, AMI and FMI in the four main cell types following the order of the UMAPs displayed above (cancer cells, CAFs, CAE and CAI cells). E. UMAPs displaying metacells assignments (dots with black borders) in each of the four cell types, unique colors indicate different metacells assignments. F. UMAP displaying cancer metacells after integration, colored by the clusters which are combination both of tissue types and patients’ treatment. G. UMAP displaying CAFs metacells after integration, colored by cell types. H. UMAP displaying CAE metacells after integration, colored by the patients’ treatment. I. UMAP displaying CAI metacells after integration, colored by cell types.

### 2.2 Data integration allows to preserve biological variability

Prior to performing data integration we set up a method able to define a space of interest comprehensively describing most of HGSTOC variability including patient-specific variability. To this aim, the identification of the set of highly variable genes (HVG) or features is crucial. Typically, such sets are computed on the whole dataset, an approach which may thereby fail to capture genes that are characterizing only a specific patient. For these reasons, focusing on the features space of the specific cell type of interest, we first computed highly variable genes (HVG) separately for every patient of the cohort in order to capture inter-individual variability and porceeded then to select their union prior to filtering by cell type-specific normalized gene dispersion values. We named these subset of genes Highly Defining Genes (HDGs). Within the space defined by HDGs, we complemented our feature selection strategy with metacells derivation [20] aimed at reducing dimensionality of the dataset and increasing robustness of the characterization of cell states along the entire phenotypic manifold. Metacells were, again, computed separately for every single patient to retain patient-specific variability. We confirmed that metacells captured patient variability (Fig. 3E), as they spread across the phenotypic manifold following the distribution observed for the respective cells they are derived from, and they also recapitulated the cell type-specific degrees of patient heterogeneity. We then exploited a previously developed variational autoencoder [21] to integrate batch effects of the datasets while preserving patient variability. To perform data integration preserving the most of the biological signal underlying both the disease and inter-patient variability, we defined the optimal label keys to be preserved in the latent space while training the model. For each major cell type, we proceeded with a twofold strategy to define the best anchors for integration. For cancer associated fibroblasts (CAFs) and CAI cells, we defined robust cell subpopulations based on previously described markers. Specifically, for CAFs we created 4 literature-based [13], [22], [23] gene signatures describing the four main CAFs subpopulations, i.e. matrix CAFs (mCAFs), inflammatory CAFs (iCAFs), vascular CAFs (vCAFs) and STAR gene expressing CAFs (star-CAFs). The manual curation of signatures was necessary since there is no robust consensus for CAFs’ typing in published studies. For CAI cells, instead, we used the recently described gene signatures [11] that defined robustly several CAI cell subtypes, i.e. B cells, T cells, plasma cells, dendritic cells, mast cells, myeloid cells. On the other hand, for cancer and CAE metacells, given the lack of well-defined markers for the definition of robust cell subtypes, we used as anchors for integration both the tissue of origin and the treatment regimen. We were able to show that after integration cancer and endothelial cells preserved their tissue/treatment related information in homogenous populations (Fig. 3F, H) while fibroblasts and immune cells preserved the specific anchor cell types (Fig.3G, I). Our results showed that by preserving patient inter-variability we were also able to achieve good data integration that allows now to characterize homogenous biologically relevant populations.

### 2.3 Cell states variations according to treatment are tissue-dependent and cell type-specific

The identification of functional cell states is one the common use cases of transcriptional atlases, especially in cancer where it is often particularly difficult to unpack cellular heterogeneity into functionally coherent cell populations given the absence of bona fide markers. Indeed, the prediction of functional cell states is a key step to identify novel targets, either emerging from each cell state in isolation or from the interaction intervening between them and their associated TME. Here, to obtain the most comprehensive characterization of cell states for HGSTOC, we identified cell states by an approach that differs from previously described procedures based on gene modules [7], [24], [25] Specifically, we pursued an unbiased annotation of cell states based on cluster stability across different degrees of clustering granularity and went on to perform gene set enrichment analysis (GSEA) to assign biological functions to the identified clusters as a functional inroad into the cell states present in our dataset (Fig. Suppl. 2A, see Methods). We applied our strategy both to integrated cancer, fibroblasts and endothelial metacells and separately for each tissue type (primary, ascites, metastasis) to capture also cell states that may only emerge in a specific tumor tissue type. Since pan cancer studies [3], [26] identified specific states for malignant cells that are shared among many different cancer types, we investigated whether they were present in our analysis. Indeed, we found the vast majority of the described pan-cancer cell states, including cell cycle, stress response, ciliated/cell movement, cellular metabolism and ECM remodeling. We also found some atlas-specific cell states related to the interaction with the immune system (Fig. 3C). Interestingly, while the majority of them were shared across the three types of tissues, cell states related to inflammatory pathways such as IFN-response and stress response were specific for primary tissue and ascites, while ECM remodeling was only found in metastatic tissue. While the majority of cancer-derived metacells showed cell states related to cell cycle and cellular metabolism, fibroblasts derived metacells were mostly related to ECM remodeling and metabolism but also angiogenetic processes (Fig. Suppl. 2B-C). Interestingly, for endothelial-derived metacells, which represents small fraction of the entire atlas, we identified several cell states related to specific immunoreactive processes. Specifically, while endothelial cells are known to be involved in immune-related processes in ovarian cancer, here we describe for the first time cell states involved in the interaction with specific immune subpopulations (T cells, B cells, neutrophils) (Fig. Suppl. 2D), demonstrating the sensitivity our strategy in resolving, through the combination of dataset aggregation, metacells derivation and data integration, also rare cell states that are hard to identify in individual datasets. We next evaluated whether some of the identified cell states were specific to the therapeutic regimens undergone by patients and either consistent across cell types or dependent on the tissue context in which they were identified. Cell states characterizing primary tissue cancer cells following chemotherapy (CHT) are mainly related to cell movement while after NACT they are mainly related to cell cycle processes. In ascites, while cell states of unknown function are present in both CHT and NACT, the latter induces also metabolism-related states and response to extracellular signals. The latter is further amplified in metastatic tissue. Of interest, and specific of metastatic tissue, is the appearance of immunoreactive cell states that characterizes the total of CHT-treated samples and to a minor extent the NACT ones (Fig. 4A). In CAE cells, NACT leads to angiogenetic cell state in all three types of tissues but in metastatic tissue they restrict the immunoreactive phenotype to a large extent to T cells and neutrophils (Fig. 4B). In contrast with the remodeling seen for cancer and endothelial cells states, patient’s treatment does not impact CAFs composition across tissue types. Indeed, cell states here are almost superimposable to the ones identified in naïve CAFs. Of note, immunoreactivity, cell cycling and angiogenetic phenotypes are maintained after treatment only during the ascites and metastatic stages of the disease (Fig. 4C). These findings overall suggest that patient’s treatment affect in a tissue-specific fashion the cell states composition of the disease, with a particularly striking remodeling of the immune-reactive phenotypes, further reinforcing the role of tumor microenvironment in sculpting tumor composition. We also investigated the cell states characterizing CAFs cell types, given the still present uncertainty on the functions associated to each subtype. We found that mCAFs were the most heterogenous subtype in terms of biological functions (in particular in primary tissue), encompassing cellular metabolism, ECM remodeling, epithelium development, cell cycle, immunoreactivity, angiogenesis and stress response. Of note, iCAFs showed a marked evolution throughout tumor progression from primary tissue to ascites to metastasis, acquiring increasingly heterogeneous functions, among which protein catabolism emerged as a possible modality for energy supply, specific for more metastatic stages of disease. vCAFs, as expected, were characterized by cell states involved in the regulation of vascular processes and smooth muscle cells development throughout tumor evolution. starCAFs are a so far poorly characterized CAFs subpopulation and represent a small fraction of this cell type. Here we could show, that while across tumor evolution they are characterized by very heterogeneous states, they present a preponderant association with metabolic processes in the primary tissue that is quickly substituted by angiogenetic functions in later metastatic stages, pointing to a highly plastic phenotype across tumor evolution (Fig. 4C). Last, CAI metacells subtypes were found not to be tissue dependent (Fig. 4D), consistently with the homogeneity of these data observed until this point. Interestingly, while adjuvant CHT did not impact immune cell composition, we observed that the NACT regimen restricted the immune cellular landscape to CD4+ Treg cells. As final step, to verify whether our strategy highlighted phenotypic features of HGSTOC that would have been masked by inter-patient variability, we mapped the cell states here identified in metacells on the original cells composing their respective metacells (Fig. 5A). We observed that the identified cell states were now scattered along the embedding space across different patients, indicating that our strategy is able to capture common cell states, while preserving the information underlying inter-patient variability.

**Fig.4:**
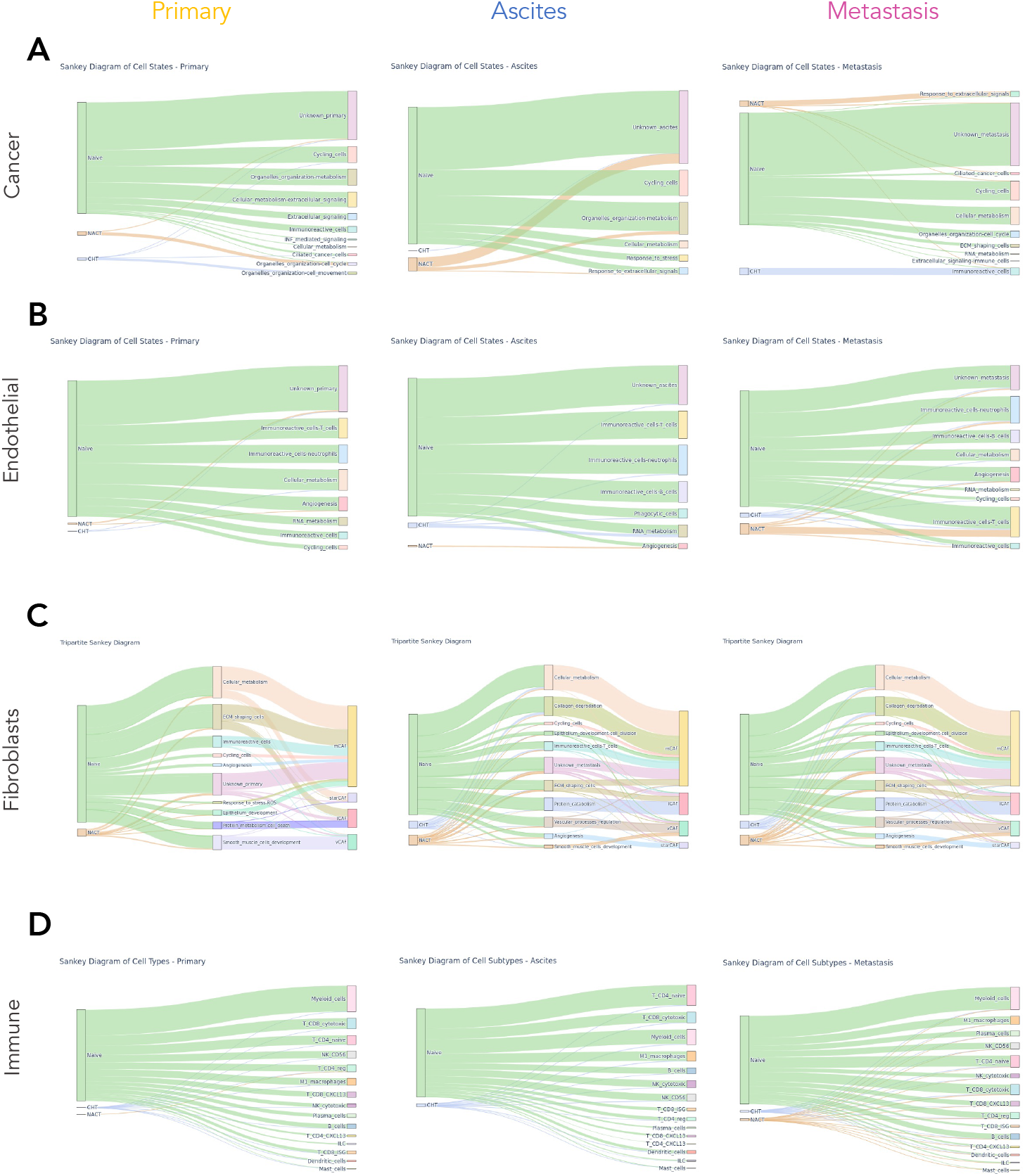
Cell states identifications across tissues and treatments. A. Sankey plot overview of the cell states annotation derived for cancer cells showing their composition within the tissue types and their abundance under different treatment conditions. B. Sankey plot overview of the cell states annotation derived for cancer-associated endothelial cells showing their composition within the tissue types and their abundance under different treatment conditions. C. Sankey plot overview of the cell states annotation derived for cancer-associated fibroblasts showing their composition within the tissue types, their abundance under different treatments and CAFs cell types. D. Sankey plot overview of the cell subtypes annotation based on gene signatures derived for cancer-associated immune cells showing their composition within the tissue types and their abundance under different treatment conditions.

**Fig.5:**
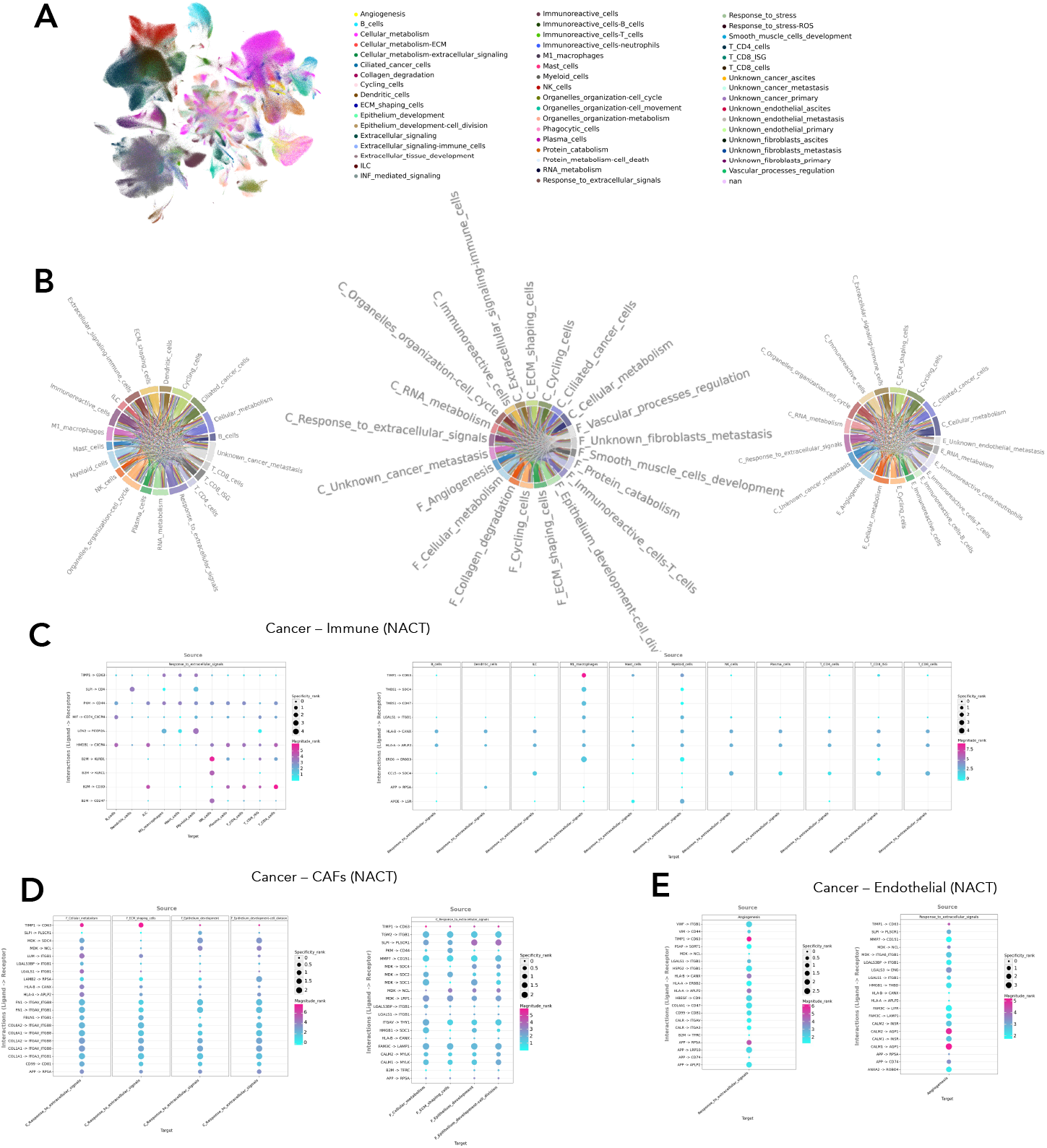
Cell to cell interaction analyses. A. UMAP displaying OvCA colored by cell states annotation derived from metacells. B. Circos plot displaying interactions between (from the left) cancer cell states and CAI cells subtypes, cancer cell states and CAFs cell states and cancer cell states and CAE cell states; each color is a cell state or cell subtype, from each of them are displayed both interactions in which they are the source and the target. C. Bubble plots showing the top 10 interactions between cancer cells population involved in “Response to extracellular signals” and CAI cells considering cancer cells as source of the ligand (on the left) and as target (on the right). Specificity rank is a measure to specifity of the interactions while magnitude rank represents the streng the of the interactions. D. Bubble plots showing the top 10 interactions between cancer cells population involved in “Response to extracellular signals” and CAFs cell states considering cancer cells as source of the ligand (on the left) and as target (on the right). E. Bubble plots showing the top 10 interactions between cancer cells population involved in “Response to extracellular signals” and CAE cell states considering cancer cells as source of the ligand (on the left) and as target (on the right).

### 2.4 CCC analyses reveals TIMP1/CD63 as a conserved interaction across disease stage following therapy

Our strategy allowed to identify cellular states with their associated biological function and to map them back to the original cells of the dataset. We thus exploited this feature to assess the presence of tumor-TME interactions in two important pathogenetic mechanisms such as metastatization and chemoresistance, evaluating in these contexts the cell states involved in specific ligand-receptor interactions. To this aim, we applied LIANA algorithm 42 to atlas subsets, concentrating on the interactions existing between cancer cells coupled with either immune cells, CAFs or endothelial cells. On each of these subsets analyses were performed based on the individual tissues (primary, ascites, metastasis) and treatments (naïve, CHT, NACT) (Fig. Suppl. 3). Given the huge amount of data and that the atlas allows very tailored analyses on the basis of the preferred topic of interest we focused on some pathogenetic aspects we found particularly interesting to look at to show the potential of this resource. To look at the effects on interactions induced by chemotherapy across tissue types we looked at conserved cell states in primary, ascites and metastasis following NACT. The only cancer cell state conserved across all tissue types was related to the response to extracellular signals. We then looked at the interaction of this cell state with cell states belonging to each major cell subpopulation. For immune cells, we showed that these cells mostly interacted with NK cells via canonical B2M/KLRD1 and B2M/KLRC1 and with T CD8 cells via B2M/CD3D suggesting that both innate and adaptive immune cells are interacting with tumor epithelial cells. In addition to this, the strongest and most specific interactions were observed with M1 macrophages and myeloid cells via TIMP1/CD63 (Fig. 5C). Considering CAFs, we found that the most conserved cell states across tissue types were the ones related to cellular metabolism, ECM shaping and epithelium development. The most interesting interactions with cancer cells existed within cellular metabolism and ECM shaping cells states via TIMP1/CD63 and LUM/ITGB1 which has been recently described as a novel prognostic marker in other types of cancer such as gastric and breast cancers[27]. Instead, cancer cells strongest interactions with these CAFs states are mostly directed toward cell states involved in the epithelium development via SLP1/PLSCR1 and MDK/NCL, this latter being already a well described interaction in ovarian cancer to be a potential prognostic and therapeutic target [28]. For endothelial cells, the angiogenetic cell state was preserved across tissues and showed interaction with cancer cells via TIMP1/CD63, APP/RPSA, HLA-A/APLP2, HLA-B/CANX. Conversely, cancer cells interacted with CAE cells via CALM2/AQP1, CALM1/AQP1 and LGALS3/ENG. Aquaporins are known to be highly expressed in ovarian cancer and to mediate pathogenetic processes such as cancer metastasis, angiogenesis and resistance to apoptosis [29], [30] while CALM2 is already been described to be involved in the same processes in other cancer types [31] but to date this is the first time that has been described their interaction in ovarian cancer. The stability of TIMP1/CD63 interactions across different stages of disease and between all cancer-associated (sender cells) and cancer cells (receiver cells) suggest that targeting TIMP1 may be a potential therapeutic strategy with broad spectrum for this disease[32].

### 2.5 CCC analyses reveals previously described tumor-TME interaction and identifies specific tissue/treatment associated ones

We next focused on the investigation of well-known interactions in ovarian cancer among ligand-receptor complexes and interactions involved in key processes related to therapeutic treatments. First, we checked the CCI between cancer and immune cells exploiting the previously defined cancer cell states to define which were the specific cancer subpopulations interacting with immune cells. The immune cell types involved in the strongest and most specific interactions with tumor cells were CD8+ T lymphocytes. The interaction was conserved independently from the disease stage and treatment of the patients (Fig. Suppl. 4). Given the more reliable prognostic role of high serum levels of LAG3 for immunotherapy treatment in OC patients, as compared to PD-1 [33] we checked for LAG3 interaction with HLA receptors and we found very robust scores both in primary tumor, ascites and metastasis and naïve and treated patients. On the contrary, we could not find other classic ligand-receptor pairs targeted by chemotherapy such as CD86/CTLA4 or PDL1/PD1 (Fig. Suppl. 4). We then performed similar analyses for the interaction of the tumor with CAFs analyzing the presence and magnitude of previously described ligand-receptor pairs interactions[28]. In particular, we found the MDK/NCL interaction predicted as the strongest and most specific one followed by MDK/LRP1 (Fig. Suppl. 5). Interestingly in naïve samples, cancer cell state related to stress response to reactive oxygen species highlighted a strong interaction with CAFs via THBS2/CD47 (Fig. Suppl. 5D). This state is lost after treatment both with CHT and NACT, consistent with its role in promoting cancer progression by remodeling of the TME [34] which should be indeed counteracted by treatment (Fig. Suppl. 5E-F). Of note, CHT treated samples show specific reduction of MDK/NCL interactions and the presence of stronger interactions of CXCL12/CXCR4 (Fig. Suppl. 5E). Overall, these results confirmed the potential therapeutic role for drugs targeting MDK [35]. For the interactions existing between cancer and CAE cells we considered both vascular and lymphatic endothelium, considering the three main ligands VEGF-A/B/C of endothelial receptors VEGFR1 (FLT1) and VEGFR2 (KDR). We scored the highest and strongest interactions between VEGF-A expressed on cancer cell states related to cellular metabolism and RNA metabolism and ITGB1 on endothelium (Fig Suppl. 6) conserved across conditions. Instead, VEGF-A strongly and specifically interacted with CD44 in CHT and NACT treated samples compared to naïve samples (Fig. Suppl. 6E, F, G). This is particularly interesting since the interaction of CD44 with VEGF-A has been described to reduce the efficacy of bevacizumab therapy in glioblastoma[36] thus suggesting a potential similar fate for ovarian cancer patient undergoing this therapeutic regimen as standard of care. Regarding the most canonical interaction of VEGF-A/B with FLT1 [37], here we predicted their interaction with NRP1, the co-receptor of FLT1, while PIGF followed the canonical direct interaction with FLT1 (Fig. Suppl. 6). This result suggests an indirect interaction of VEGF-A/B with their receptor but also that the angiogenesis pathway is active since without the interaction with the co-receptor this would have been inhibited [38]. Regarding the interaction with the lymphangiogenesis pathway, we were able to score the interaction of VEGF-C with LYVE1 and NRP2 which are specific receptors of lymphatic endothelium [39]. To summarize, these results indicate the reliability of the atlas to retrieve well-known pathogenetic molecular processes but also shows the possibility to identify new therapeutic targets to be exploited for further investigation in the laboratory and clinical settings.

### 2.6 SCAIP allows expansion of the OvCA

One of the defining features of atlases is the possibility to iteratively expand it as new relevant data are generated (a process known as out of sample extension). This process allows to progressively increase the robustness of representation for the considered disease, especially for less represented populations and categories of metadata (e.g. treated samples). To test the possibility to extend OvCA we downloaded a new dataset [40] of 10 HGSTOC patients including samples from all three tissue types. Exploiting the reference model generated with the training on OvCA data we mapped the query dataset on the atlas reference. Reference mapping in latent space allowed to integrate the query within atlas data (Fig. 6A). We then tested whether our model was able to correctly map the new metacells on the embedding space positions corresponding to the categories they belonged to in the atlas (Fig. 6B) and quantitatively evaluated the accuracy of mapping of the new dataset by performing gaussian kernel density estimates (KDE). We could score 98.12% accuracy for primary cells, 79.55% for ascites cells and 98.71% for metastasis derived cells (Fig. 6C-D), suggesting that our model is able to reliably perform out of sample extension tasks and will allow to iteratively expand our atlas with newly generated datasets. Finally, we were able to transfer our identified cell states to the newly added dataset, confirming the robustness of out of sample extension algorithm (Fig. 6E).

**Fig.6:**
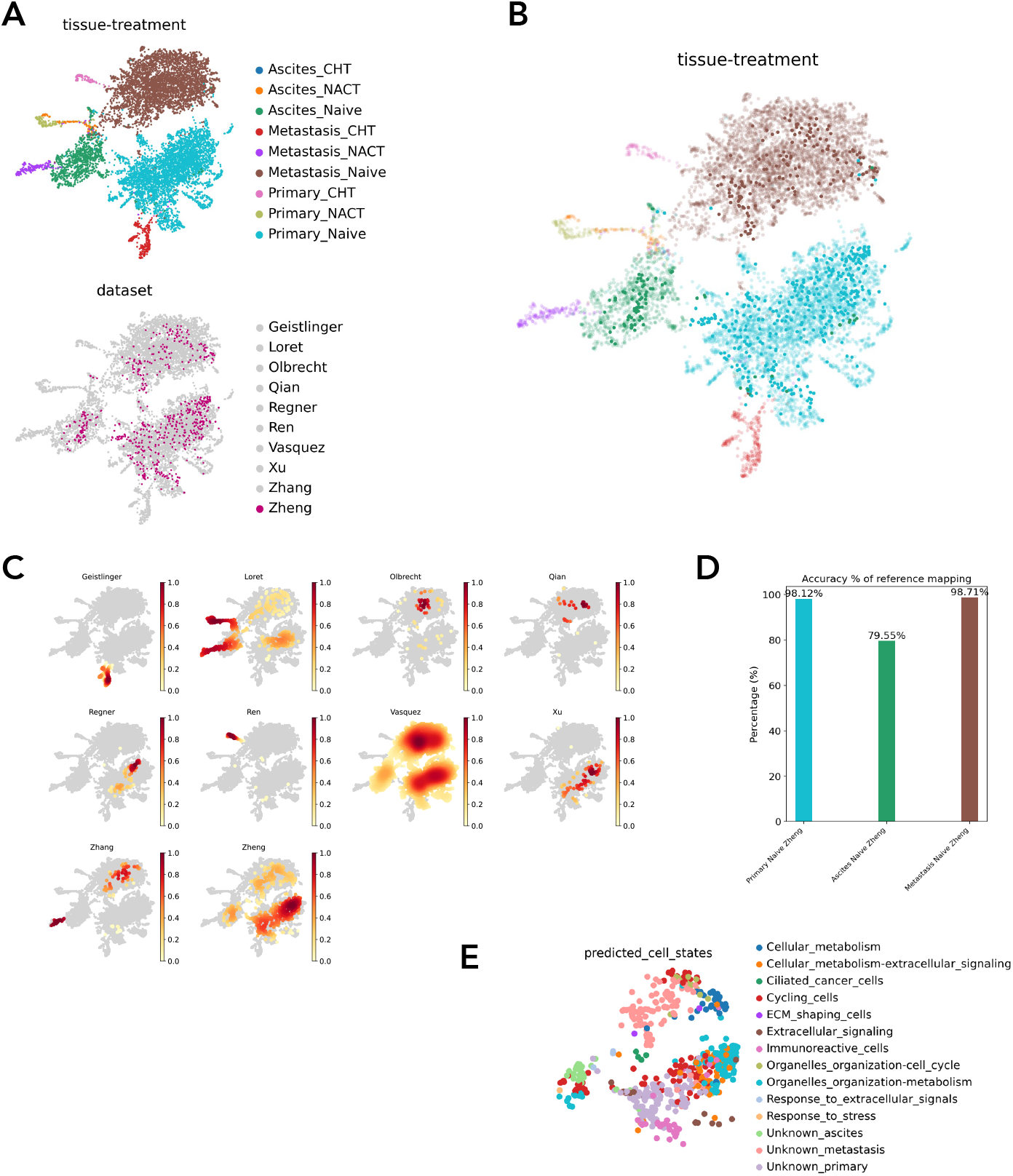
Out of sample extension. A. UMAPs built on latent space embeddings of OvCA cancer metacells integration with Zheng derived cancer metacells, colored by the cluster’s combination both of tissue types and patients’ treatment (upper plot) and by dataset (lower plot) where OvCA datasets are displayed in grey while Zheng in fuchsia. B. UMAPs built on latent space embeddings of OvCA cancer metacells integration with Zheng derived cancer metacells, colored by the clusters which are combination both of tissue types and patients’ treatment, here transparency is added on OvCA metacells while full color is applied to Zheng derived metacells. C. Density plots displaying the contribution of the different datasets to metacells distribution within extended OvCA. D. Percentages of accuracy of assignment of Zheng metacells to the clusters combination both of tissue types and patients’ treatment with respect to their original metadata annotations. E. UMAPs built on latent space embeddings of OvCA cancer metacells integration with Zheng derived cancer metacells, displaying only Zheng derived cancer metacells and colored by label trasnferred from OvCA.

## 3 Discussion

In this study we present a robust strategy for the integration of highly heterogeneous single cell transcriptomic datasets that allows to identify disease-associated cell states while preserving inter-sample biological variability. We show how this method can be successfully applied to a paradigm of cancer heterogeneity, generating the first HGSTOC single cell reference atlas (OvCA). By integrating samples from 79 different patients from the major studies on HGSTOC to date, OvCA provides a comprehensive description of the cellular and molecular diversity of this disease, offering a reference resource that can be easily accessed and explored by the scientific community.

Thanks to the integration strategy and feature selection associated with the atlas (SCAIP), we were able to describe the cell states characterizing the heterogeneity of the main populations making up this cancer ecosystem (cancer cells and their associated normal cells, encompassing fibroblasts, immune and endothelial cells) and associate them with specific disease stages and treatments (from primary tumor to ascites to metastasis). Also, we evaluated the ability of the atlas to capture the evolving landscape of cell-to-cell interactions, concentrating on the identified cell states in different major cell subtypes and the prediction of intervening relationships between them. The analyses we detailed are only a small representation of the possibilities that OvCA affords. Also, we built this resource envisioning the possibility of an iterative expansion of the atlas (out of sample extension) thanks to the model provided for reference mapping that will allow to update the atlas through version releases as soon as new datasets will be generated. This aspect is crucial since one inherent limitation of this first version of OvCA, as of cancer atlases in general, is the reduced representation of post-treatment samples. Also, underrepresented or functionally undefined cell states would benefit in their characterization from the addition of newly added data. This process will not only be limited to single cell transcriptomics data. Indeed, recent label transfer strategies will allow to integrate also additional layers of information, such as the ones derived by chromatin accessibility data, protein phenotyping and spatial transcriptomics data [41], de facto creating a multilayered cartography of disease complexity. As OvCA seeds a fundamental resource poised to evolve with the community over time, the companion implementation of an open access web resource based on CELLxGENE (https://cellxgene.cziscience.com/) provides not only direct access to our findings but also a flexible tool to address users’ own scientific questions through data exploration and analysis. Together, all these features make OvCA an accelerator of HGSTOC research towards the identification of novel therapeutic targets. Finally, beyond HGSTOC, we envision the methodologies we developed for the generation of this resource (patient-specific identification of HVG and metacell computation, alongside SCAIP) to be widely exploited, enabling an integration and characterization of heterogeneous single cell transcriptomic data that can be easily translated to other settings as foundational resources for precision medicine across disease domains.

## 4 Glossary

Cell types: groups of cells defined based on literature-derived gene expression signatures
Cell states: groups of cells defined based on the biological features derived from the functional annotation of differentially expressed gene modules
Out of sample extension: process that exploits a previously generated model to integrate new datasets into the atlas
Functional annotation: for ease of navigation to the final user of the CELLxGENE platform, functional annotation comprises both the cell states identified in the manuscript and the cell subtypes related to cancer associated immune cells since the latter are superimposable to their associated biological function
CAI: cancer-associated immune
CAF: cancer-associated fibroblast
CAE: cancer-associated endothelial

## 5 Data and code availability

Individual single cell datasets included in the atlas can be found at the data availability sections of their respective published papers. A section of our code will be dedicated to download these datasets from our server (https://repo.bioserver.ieo.it/GT/). The code to reproduce the atlas, including data integration strategy, can be found at https://github.com/GiuseppeTestaLab/atlas_project. The atlas resource to be used for data exploration can be found at https://cellxgene.bioserver.ieo.it

